# A comprehensive catalog with 100 million genes and 3,000 metagenome-assembled genomes from global cold seep sediments

**DOI:** 10.1101/2023.04.10.536201

**Authors:** Yingchun Han, Chuwen Zhang, Zhuoming Zhao, Yongyi Peng, Jing Liao, Qiuyun Jiang, Qing Liu, Zongze Shao, Xiyang Dong

**Affiliations:** Key Laboratory of Marine Genetic Resources, Third Institute of Oceanography, Ministry of Natural Resources, Xiamen 361005, China; School of Marine Sciences, Sun Yat-Sen University, Zhuhai 519082, China

**Author notes:** Correspondence can be addressed to Xiyang Dong.

## Abstract

Cold seeps harbor abundant and diverse microbes that represent a tremendous potential for biological applications and also have a significant influence on biogeochemical cycles. Though recent metagenomic studies have expanded our understanding of the microbial community and function of seep microorganisms, the knowledge of diversity and genetic repertoire of global seep microbes is lacking. Here, we collected a compilation of 165 metagenomic data from 16 cold seep sites across the globe to construct comprehensive gene and genome catalogs. The non-redundant gene catalog was comprised of 147 million genes (clustered at 95% amino acid identity), and 35.72% of them could not be assigned to a function with the currently available databases. A total of 3,164 species-level representative metagenome-assembled genomes (MAGs) are obtained, most of which (94.31%) belong to novel species. Of them, 81 ANME species are identified covering all subclades except ANME-2d, and 23 syntrophic SRB species spanning Seep-SRB1a Seep-SRB1g, and Seep-SRB2 clades. The non-redundant gene and MAGs catalogs are a valuable resource that enables expanded knowledge of the structure and functions of cold seep microbiomes.

## Background & Summary

Cold seeps occur in continental margins worldwide. At these sites, methane-rich fluids migrate from deep subsurface to the sediment-water interface^1^. Methane is a climate-active greenhouse gas approximately 30 times more potent than carbon dioxide^2^. Fortunately, in seep sediments, methane could be consumed through the process of anaerobic oxidation of methane (AOM). This process removes roughly 90% of the methane produced globally in marine sediments, acting as an efficient methane filter^3,4^. As a consequence, these seeps are critical in regulating the amount of methane released into the overlying waters and atmosphere, and they play a vital role in mitigating global warming. AOM is performed by anaerobic methanotrophic archaea (ANME). Normally ANME rely on a syntrophic partner to couple CH_4_ oxidation to the reduction of terminal electron acceptors, such as sulfate, iron, nitrate, and manganese^5,6^. AOM coupled to sulfate reduction is the primary biological process in seep sediments since sulfate is the dominant anion present at the marine sediment-water interface. High rates of AOM fueled by near-saturated methane concentrations would rapidly consume sediment pools of any individual electron acceptor, creating unique geobiological engines that contribute significantly to local and global biogeochemical cycles^1^.

Cold seeps are deep-sea oases that support immense biodiversity and where specialization and adaptation create extraordinary lifestyles^1^. However, the majority of microorganisms found in seeps are not characterized yet^7^. Culture-independent metagenomic techniques are the key to unraveling genetic diversity and metabolic potential of uncharacterized microbes, and have been applied to identify thousands of microorganisms and their metabolic versatility. Recently, the microbial community and function of cold seep sediments are increasingly studied with metagenomes obtained from different sea areas^7-10^. However, there are no large-scale gene and genome catalogs available for the microbiome of global cold seeps. A comprehensive gene catalog of cold seeps can serve as a reference for mining novel genetic resources in the deep sea, such as biosynthetic gene clusters (BGCs), which will also boost downstream biomedicine industries.

ANME and their syntrophic SRB partners play a crucial role in the regulation of both carbon and sulfur cycles of seeps. Through their mutualistic interactions, they perform AOM, leading to the reduction of methane release and the generation of inorganic carbon and sulfide. These processes are of significant importance for both local and global biogeochemical cycles, underscoring the essential role of these microorganisms in deep-sea ecosystems. Although previous findings have revealed various lineages of ANME and SRB in seep sediments^11-13^, there is currently a lack of a comprehensive genome catalog of these lineages in cold seep sediments globally. High-quality and extensive reference genomes of global seep microbiome can improve the resolution and accuracy of taxonomic and functional analyses, and provide the opportunity for largescale comparative genomes^14-17^, especially for elucidating the physiological basis of ANME-SRB interactions.

Here, we collected metagenomic sequence data from 165 sediment samples at 16 cold seeps across the Pacific, Atlantic, and Arctic Oceans **(Fig. 1)**, encompassing gas hydrates (n = 4), methane seep (n = 14), oil and gas seeps (n = 4), mud volcanoes (n = 2) and asphalt volcanoes (n = 1). Sediment samples span different depths and redox conditions, from the oxic sediment-water interface into anoxic layers down to 68.55 m below the sea floor (mbsf) **(Supplementary Table 1)**. Metagenomic raw reads were quality-controlled, generating 9.5 Tb of clean reads. All clean reads were assembled, resulting in a total of 573,669,419 contigs. Then we selected assembled contigs with a length greater than 500 bp for predicting protein-coding sequences (CDS), and obtained 373,051,862 CDSs in total. After clustering these sequences at 95% amino acid identity^15,16^, we obtained 147,289,169 protein clusters, forming the non-redundant gene catalog **(Fig. 1c)**. The catalog is the most comprehensive gene catalog of the seep microbiome up to data. And compared to microorganisms from other habitats, the microbes found in cold seeps possess a relatively high number of non-redundant genes. For example, Liu et al. ^15^ sequenced 883 bacterial genomes that were cultured and 85 metagenomes from 21 Tibetan glaciers covering diverse habitats, and constructed a gene catalog containing 25,320,330 genes. Zeng et al. ^16^ specifically analyzed 6,122 fecal metagenomes from children under the age of three and generated the Early-Life Gut Proteins (ELGP) catalogs, including 4,036,936 gene clusters.

**Figure 1.**
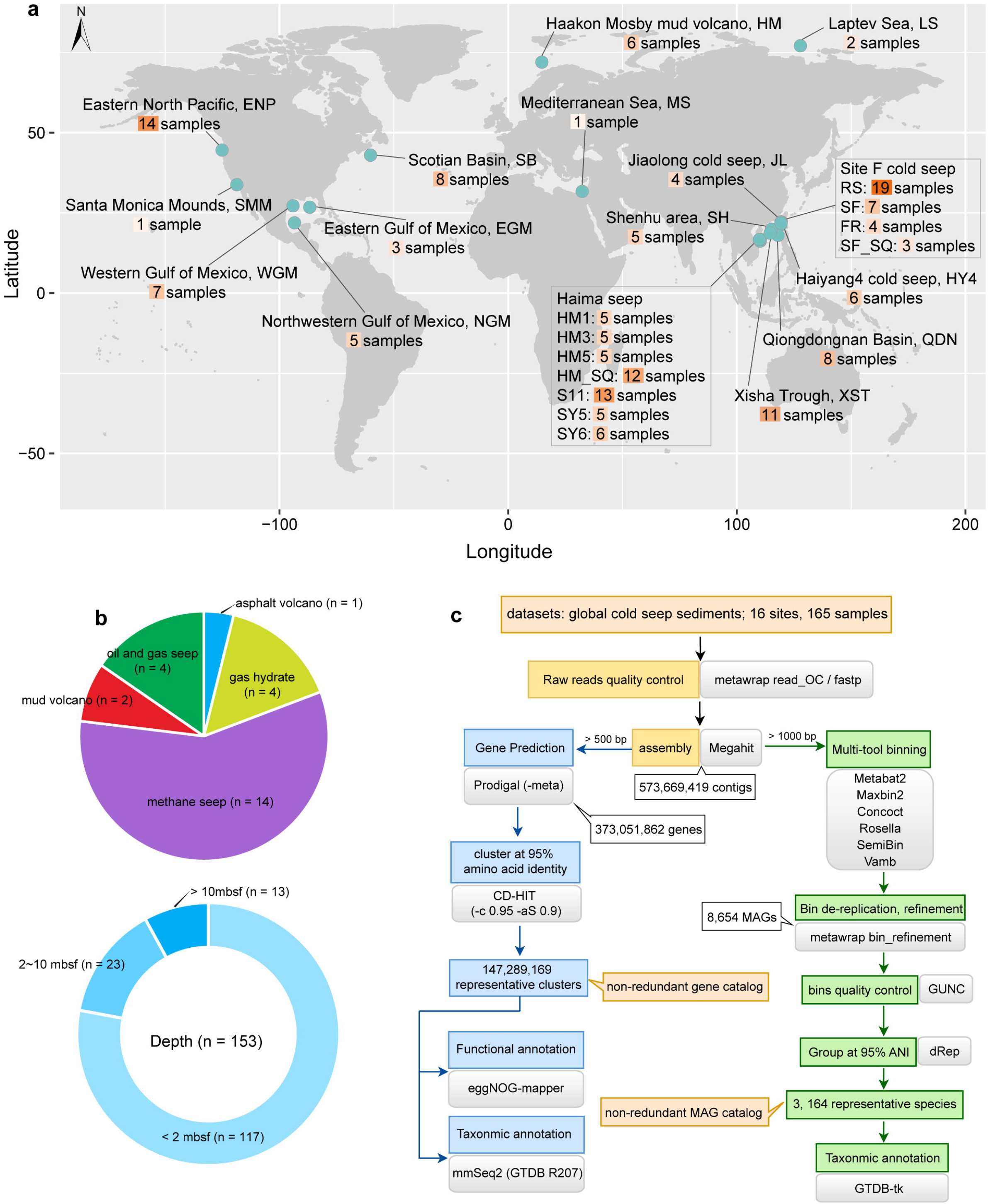
Overview of the studied areas and bioinformatics workflow. (a) Geographic distribution of the 16 global cold seep sites where metagenomic sequencing data were collected. The map was drawn using maptools and ggplot2 packages in R v4.0.3. (b) Numbers and proportions of cold seep samples classified according to their types and depths. (c) Overview of the computational pipeline to generate the nonredundant gene and MAG catalogs.

To elucidate the functional diversity of cold seep microbes, we used the available databases, including eggNOG, Pfam, KEGG, level-4 Enzyme Commission categories (EC), Gene Ontology (GO), and CAZy, to annotate the functions of the non-redundant genes. We found that 64.28% of the non-redundant genes matched to at least one of the databases of eggNOG (n = 88,929,242; ∼60%), Pfam (n = 85,404,569; ∼58%), KEGG (n = 48,756,524; ∼33%), EC (n = 27,619,712; ∼19%), GO (n = 5,966,227; ∼4%) and CAZy (n = 1,514,988; ∼1%) **(Fig. 2)**. After analyzing the annotated genes based on the eggNOG database **(Fig. 2c)**, the predominated category was “Function unknown” (n = 17,018,774). This category includes proteins that have not yet been characterized or to which there is insufficient information to assign a specific function. The proportion of unannotated genes was 35.7% (n = 52,606,290; **Fig. 2a)**, larger than that of the nonredundant global microbial gene catalog (∼27%)^14^, the Tibetan glacier gene catalog (22%)^15^, the early-life human gut microbial protein catalog (30%)^16^, and the nonredundant ruminant gastrointestinal tract microbial gene catalog (34%)^18^. These results indicate that nearly half of the genes of cold seep microorganisms were so far functionally unidentified, suggesting that cold seeps harbor numerous unknown functional genes. We used the Genome Taxonomy Database (GTDB R207)^19^ to annotate the taxonomy of the non-redundant amino acid sequences. A notable percentage of non-redundant sequences (n = 44,441,531; ∼30%,) could not be classified as belonging to any microorganisms in GTDB, suggesting that these sequences may be attributed to novel prokaryotic microbes **(Fig. 2d)**. Around 9% (n = 13,154,825) of nonredundant sequences could only be identified as either bacteria or archaea, and could not be further classified at the Phylum level **(Fig. 2d)**. The results of taxonomic classification further confirm that this gene catalog contained lots of untapped genetic resources. In summary, the non-redundant gene catalog is an unprecedented-scale gene resource of seep microbes and can be used as a reference for future studies on mining novel bioresources from deep-sea cold seep ecosystems.

**Figure 2.**
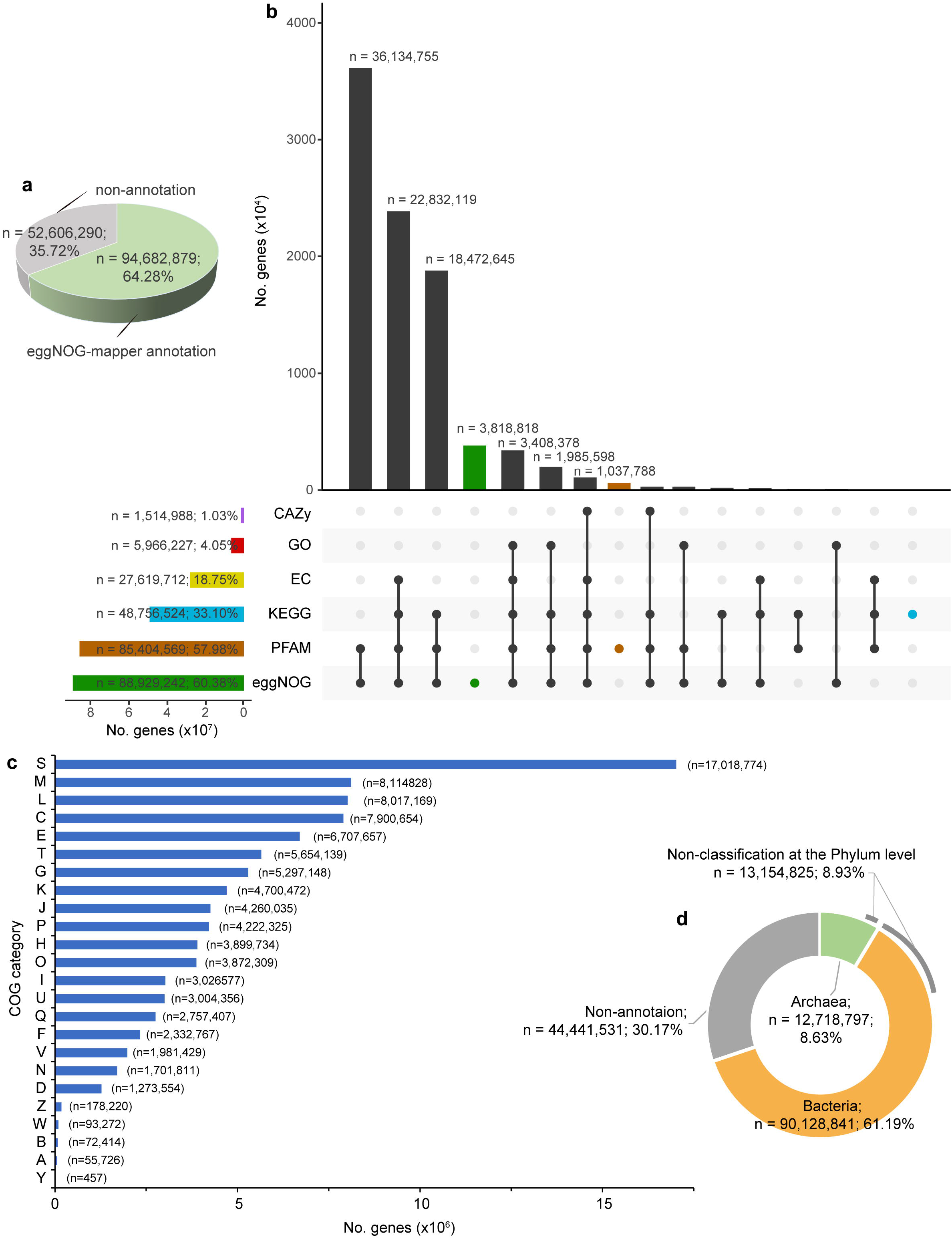
Functional and taxonomic characterization of non-redundant gene catalog. (a) An overview of annotations for the non-redundant gene catalog. Non-annotation indicates that these genes were not annotated in at least one of the databases of eggNOG, Pfam, KEGG, EC, GO and CAZy. (b) Number of genes with functional annotations across the six functional databases. Vertical bars represent the number of genes unique (color) to each functional database or shared (black) between different functional databases. Horizontal bars in the lower panel indicate the total number of genes with functional annotations in each database. (c) Functional annotations at the COG category level. S: Functional unknown. (d) Breakdown of taxonomic annotations for the non-redundant gene catalog.

A total of 8,634 MAGs were recovered, which met or exceeded the medium-quality (≥ 50% completeness and <10% contamination) based on the “Minimum information about a metagenome-assembled genome (MIMAG)” standard^20^. To control the quality of the genome more strictly, we selected those genomes that were free of chimerism (passed by GUNC^21^), resulting in 7,776 MAGs. All MAGs were clustered using a 95% whole-genome average nucleotide identity (ANI) cut-off^22,23^, defining 3,164 specieslevel clusters. The 3,164 species clusters covered various prokaryotic lineages spanning 113 phyla (97 bacterial and 16 archaeal) based on taxonomic annotations from the GTDB Database (R207)^19,24^. They were highly represented by bacterial phyla of Chloroflexota (n = 371), Proteobacteria (n = 335), Desulfobacterota (n = 306), Planctomycetota (n = 190), Patescibacteria (n = 152) and Bacteroidota (n = 151), and archaeal phyla of Halobacteriota (n = 129), Thermoplasmatota (n = 108), Thermoproteota (n = 98), Asgardarchaeota (n = 95) and Nanoarchaeota (n = 47) **(Fig. 3b)**. According to the results of taxonomic classification, four species clusters (CSMAG_1466: completeness 98.9% contamination 0; CSMAG_2247: com 60.83% con 1.19%; CSMAG_2329: com 62.74% con 1.59%; CSMAG_3128: com 85.47% con 3.75%) were not assigned to any existing phylum, suggesting that these species potentially belong to new phyla. Furthermore, 44 classes, 184 orders, 412 families, 1,043 genera and 2,984 species do not have representatives in the GTDB **(Fig. 3b)**, representing potential novel lineages. Overall, ∼94% of representative species show substantial novelty, which exhibit low sequence identities to existing genomes in the database, suggesting that cold seep harbor a diverse range of novel microbial species. The non-redundant MAG catalog represents an unparalleled genome resource, considerably expanding the phylogenetic diversity of the cold seep microbiome.

**Figure 3.**
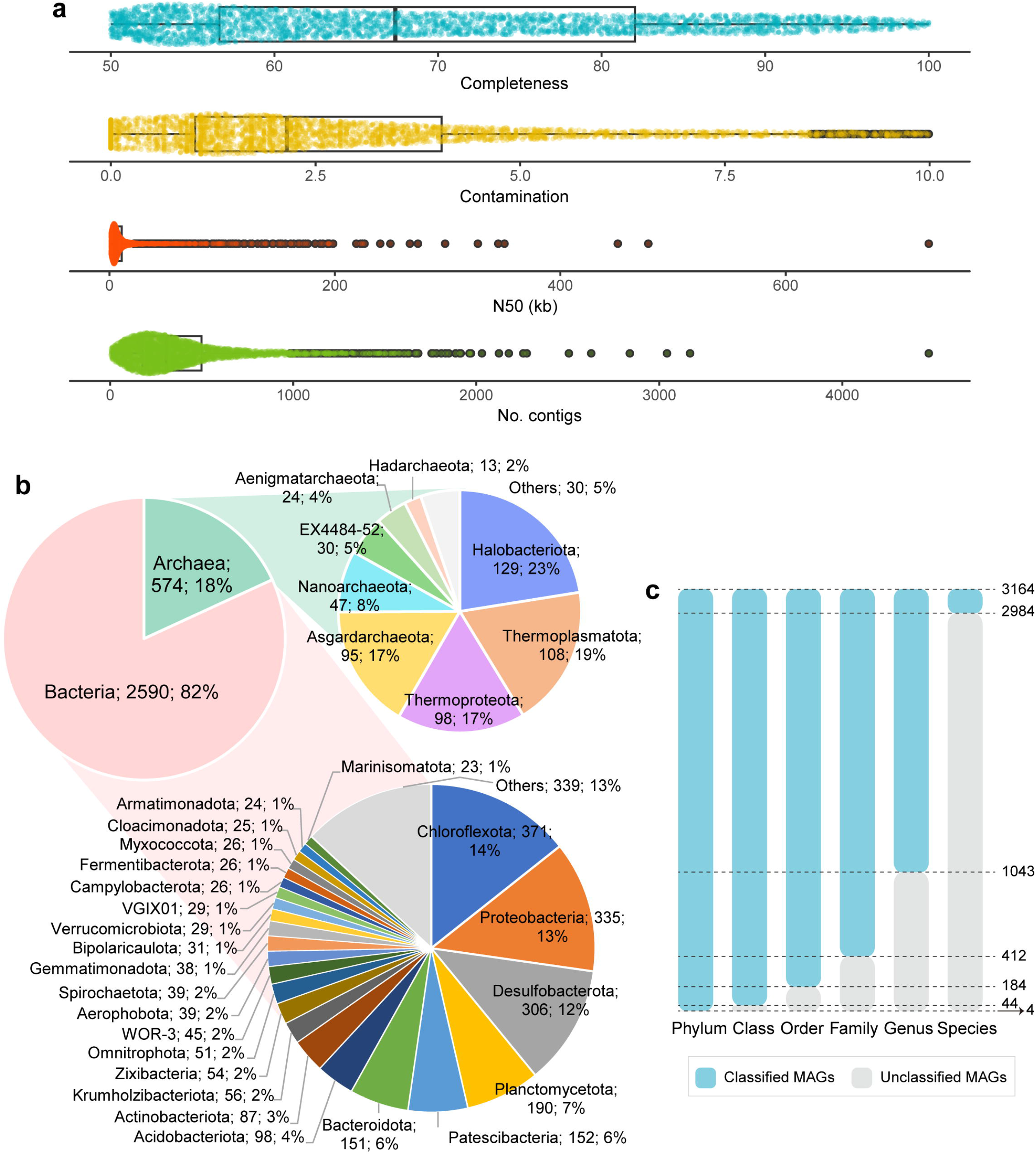
Quality, and novelty of non-redundant MAGs. (a) Genome statistics for the representative species of non-redundant MAGs. (b) Taxonomic classification (domain and phylum levels) of the species-level representative MAGs. (c) Taxonomic novelty of the representative species.

To identify ANME and syntrophic SRB lineages in seeps, we constructed phylogenomic trees of ANME and syntrophic SRB based on archaeal and bacterial single-copy marker genes, respectively. We collected 41 previously published ANME genomes^25-33^, including all of the currently described subclades (ANME-1, ANME-2a, ANME-2b, ANME-2c, ANME-2d, and ANME-3), as well as 60 SRB genomes, including syntrophic (HotSeep-1, Seep-SRB2, Seep-SRB1a, and Seep-SRB1g) and non-syntrophic SRB^34-37^. According to the phylogenomic distance, we found 81 ANME genomes spanning five subclades including ANME-1 (n = 38), ANME-2a (n = 16), ANME-2b (n = 1), ANME-2c (n = 24), and ANME-3 (n = 2) **(Fig. 4)**. Recently, Chen et al. ^13^ obtained 63 ANME MAGs from global methane seeps, including 50 MAGs collected from publicly available databases and 13 MAGs recovered from the Haima methane seep. After dereplication, 47 MAGs were retained, and these ANME MAGs were distinctly clustered into three clades, including ANME-1a/b (n = 21), ANME-2a/b (n = 11), and ANME-2c (n = 15)^13^. These ANMEs were obtained only from the type of methane seeps, and their quantities were smaller than those recovered in this study. We also identified 23 syntrophic SRB genomes **(Fig. 5)** spanning three clades (Seep-SRB2, n = 8; Seep-SRB1a, n =14, and Seep-SRB1g, n = 1). The compendium of ANME and syntrophic SRB will contribute to expanding the understanding of physiological basis of ANME-SRB interactions.

**Figure 4.**
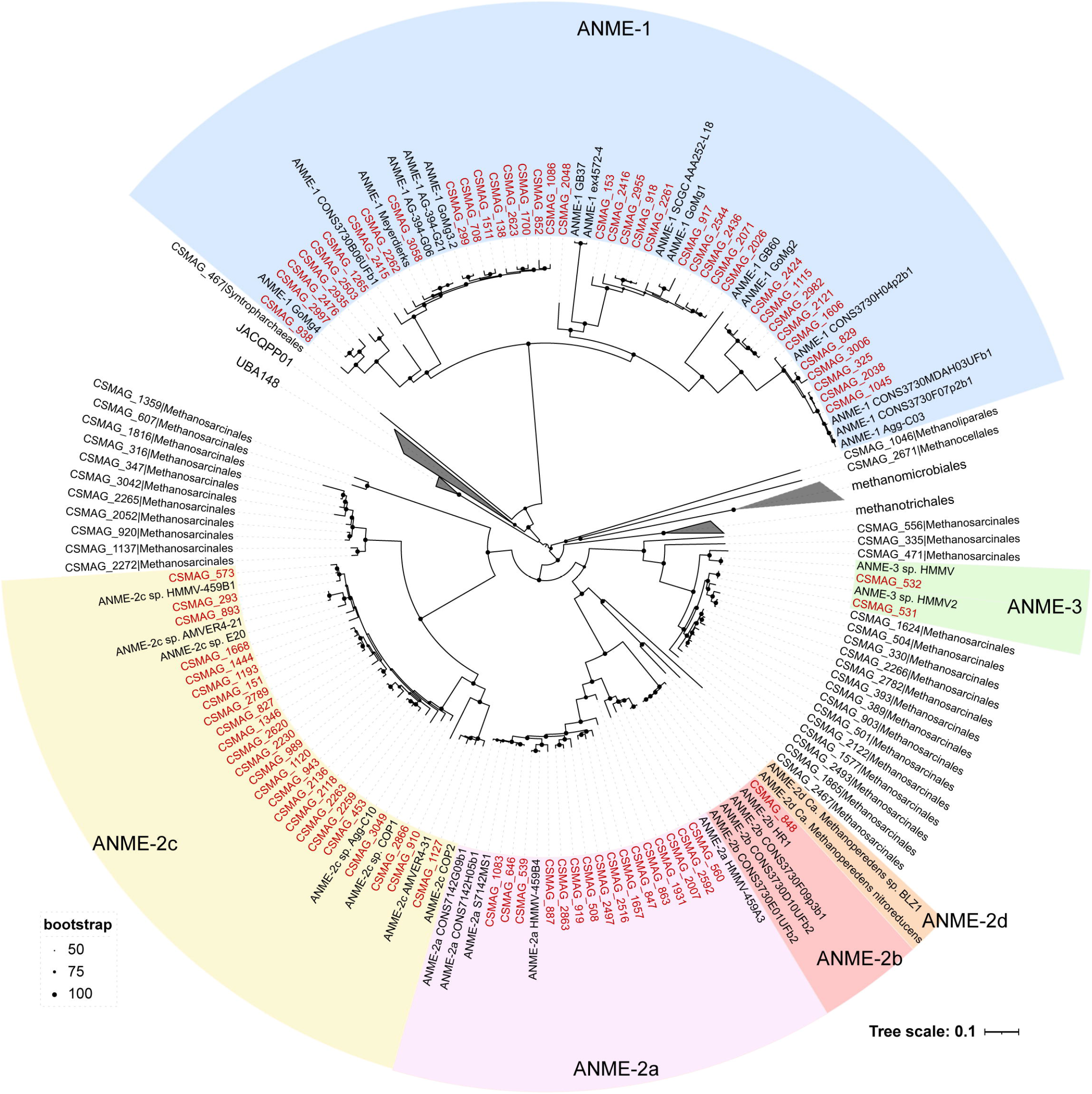
Phylogenomic tree of ANME genomes and related archaea. The phylogenomic tree was constructed from 41 previously published ANME genomes and 135 MAGs belonging to Halobacteriota from this study. The tree was constructed by maximum likelihood method using a concatenated alignment of 53 conserved archaeal single-copy marker genes.

**Figure 5.**
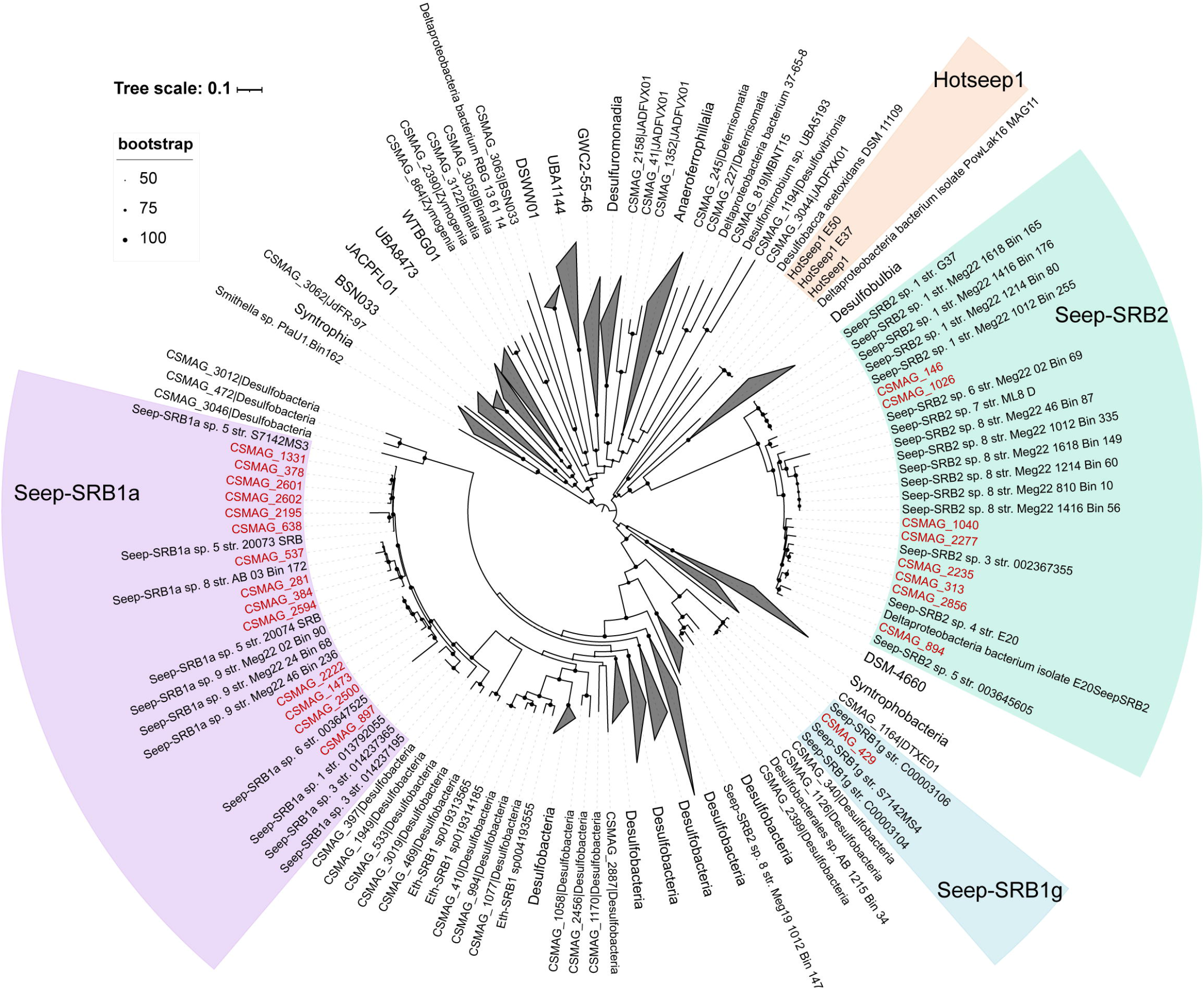
Phylogenomic tree of syntrophic SRB genomes. The tree was constructed that included 60 reference SRB genomes collected from previous studies and 327 MAGs assigned to Desulfobacterota from this study. The tree was constructed by the maximum likelihood method using a concatenated alignment of 120 conserved bacterial single-copy marker genes.

## Methods

### Collection of metagenomes

Metagenomic data sets were comprised of 165 sediment samples (0 to 68.55 mbsf) collected from 16 globally distributed cold seep sites **(Fig.1a; Supplementary Table 1)**. These sites are as follows: Eastern North Pacific (ENP), Santa Monica Mounds (SMM), Western Gulf of Mexico (WGM), Eastern Gulf of Mexico (EGM), Northwestern Gulf of Mexico (NGM), Scotian Basin (SB), Haakon Mosby mud volcano (HM), Mediterranean Sea (MS), Laptev Sea (LS), Jiaolong cold seep (JL), Shenhu area (SH), Haiyang4 (HY4), Qiongdongnan Basin (QDN), Xisha Trough (XST), Haima seep (HM1, HM3, HM5, HM_SQ, S11, SY5, SY6) and site F cold seep (RS, SF, FR, SF_SQ). Paired-end sequencing data from ENP, SMM, WGM, NGM, HM, MS, LS and part of site F (RS and FR) were downloaded from the National Center for Biotechnology Information-Sequence Read Archive (NCBI-SRA) and European Bioinformatics Institute-European Nucleotide Archive (EBI-ENA) according to the accession numbers published in each study^8-10,35,38-41^. The remaining 106 metagenomic datasets used in this study were obtained from our previous publications^7,12,42-49^. Detailed sequencing information is available in **Supplementary Table 1**. These metagenomic samples were collected from a range of cold seeps, including oil and gas seeps, methane seeps, gas hydrates, asphalt volcanoes, and mud volcanoes. The samples were taken from various depths and under different redox conditions, from the oxic sediment-water interface and extending down to anoxic layers as deep as 68.55 meters below the sea floor.

### Contig assembly, gene prediction and gene catalog construction

Metagenomic sequence data were quality controlled using Read_QC module (--skip-bmtagger) within the metaWRAP v1.3.2 pipeline^50^ and fastp v0.23.2^51^ (default parameters). After quality control, 9.5 Tb of clean reads were remained for subsequent analyses. Clean reads from each cold seep sediment at individual depths were assembled using MEGAHIT (v1.1.3 and v1.2.9)^52^ with default parameters, and coassemblies were performed by combining metagenomes from all depths of each cold seep sediment using MEGAHIT v1.1.3 ^52^ with the options “--k-min 27 --kmin-1pass -- presets meta-large”. Detailed assembly statistics were summarized in **Supplementary Table 1**. Assembled contigs (length > 500 bp, n = 225,026,054) were used to predict CDS with Prodigal (v2.6.3; parameter: -meta)^53^, which generated 373,051,862 protein sequences. These sequences were then clustered at 95% amino acid identity using CD-HIT (v 4.8.1)^54^ with the parameters: “-c 0.95 -aS 0.9 -g 1 -d 0”. This resulted in a nonredundant gene catalog comprising 147,289,169 representative clusters.

### Functional and taxonomic annotations of the non-redundant gene catalog

The representative amino acid sequences from each cluster were functionally annotated using eggNOG-mapper v2.1.9^55,56^ with default parameters. The functional annotations including eggNOG 5.0, Pfam 33.1, KEGG, EC, GO, and CAZy were derived from the eggNOG-mapper results. MMseqs2 (v13.45111) taxonomy^57^, with parameter “--tax-lineage 1”, was used to assign taxonomic labels to each representative amino acid sequence. GTDB R207 was used as the reference database^19^. The MMseqs2 taxonomy uses an approximate 2bLCA (lowest common ancestor, LCA) approach.

### Metagenomic binning and non-redundant MAG catalog construction

Assembled contigs were filtered by length (> 1000 bp) for subsequent binning. Filtered assembled contigs were binned using metaWRAP v1.3.2^50^ binning module (parameter s: -metabat2, -maxbin2, -concoct, -universal), SemiBin v1.4.0 with single_easy_bin mode (default parameters)^58^, and Rosella v0.4.1 (default parameters) (https://github.com/rhysnewell/rosella). For S11 and RS, individual assemblies from these samples were concatenated and binned separately using the VAMB tool (v3.0.2; --minfasta 200000 -o C)^59^. Afterwards, the produced bins from each binning tool were integrated and refined using the Bin_refinement module of metaWRAP v1.3.2 pipeline^50^ with settings “-c 50 -x 10”. The completeness and contamination of refined bins were evaluated with CheckM v1.2.1^60^. Then, the resulting 8,654 MAGs were checked by GUNC v1.0.5^21^ (default parameters) to remove genomes potentially containing chimerism based on “pass.GUNC”. All MAGs were dereplicated at species level using dRep v3.4.0 (parameters: -comp 50 -con 10)^61^ with an average nucleotide identity (ANI) cutoff value of 95%. Representative genomes were selected based on the dRep scores derived from genome completeness, contamination and assembly N50. A total of 3,164 MAGs with the highest dRep score from each species cluster were selected as the representative species.

### Taxonomic annotations and novelty evaluation of MAGs

Taxonomic annotations of each MAG were performed using GTDB-Tk^24,62^ toolkit v2.1.1 with the “classify_wf” workflow (default parameters) against the reference database (R207). Afterwards, we used classification results based on GTDB R207 to evaluate the novelty of each MAGs^15,63^. The novelty of MAGs was assessed through various factors, including their placement in the GTDB reference tree, RED values and ANI in comparison to reference genomes, as determined by GTDB-Tk^24,62^. Briefly, genome classifications were primarily determined by their placement in the GTDB reference tree. If the placement of a genome is ambiguous, the relative evolutionary divergence (RED) value is used for further classification based on well-established standards for each taxonomy level in GTDB. And the assignment of a genome to an existing species was based on the ANI value to reference genomes with a threshold of 95%.

### Genomes for ANME and their syntrophic SRB

To explore the diversity of ANME lineages in global cold seep sediments, a phylogenomic tree was constructed that included 41 previously published ANME genomes^25-33^ and 135 MAGs belonging to Halobacteriota from this study. These published ANME genomes cover all of the currently described subclades: ANME-1, ANME-2a, ANME-2b, ANME-2c, ANME-2d, and ANME-3. To identify their syntrophic SRB, we constructed a phylogenomic tree of concatenated marker genes from 60 reference SRB genomes^34-37^ (including syntrophic SRB: HotSeep-1, Seep-SRB2, Seep-SRB1a and Seep-SRB1g, and non-syntrophic SRB) and 327 MAGs assigned to Desulfobacterota from this study. The concatenated multiple sequence alignment of genomes based on 53 archaeal and 120 bacterial single-copy marker genes was produced via the identify and align workflow of GTDB-Tk (v2.1.0)^24^. The maximum likelihood tree was constructed using IQ-TREE (v2.2.0.3)^64^ with the “-m MFP -B 1000” options. All produced phylogenomic trees were visualized using iTOL v6 ^65^.

### Data Records

All raw data analyzed in this study are publicly available. Supplementary Table 1 contains the accession numbers for all the metagenomes used. The non-redundant gene catalog and files for the phylogenomic trees are available on figshare (10.6084/m9.figshare.22568107). Genome sequences of non-redundant MAGs have been uploaded to figshare (10.6084/m9.figshare.22568107) and submitted to NCBI under BioProject accession number PRJNA950938.

### Technical Validation

To maximize the number of genes and ensure the quality of genes, we selected assembled contigs with a length greater than 500 bp to predict CDS as suggested by previous studies^15,18,66^. Then we selected assembled contigs by length (> 1000 bp) for metagenomic binning. The number of metagenomic samples collected from S11 (n= 13) and RS (n=19) was relatively larger than those obtained from other sites, making it challenging to bin the co-assemblies of the samples from these sites in terms of computation resources. Thus, individual assemblies from the S11 and RS sites were concatenated and binned separately using the VAMB tool. And the quality of MAGs was strictly controlled according to the following standards: 1) completeness > 50% & contamination < 10%; 2) genome sequences without chimerism (details in **Supplementary Table 2**).

### Usage Notes

The datasets analyzed in this study are the largest cold seep metagenomic datasets considered to date. Researchers could use the gene catalog of seeps to compare genes of interest to other habitats, such as glaciers, polar regions and hydrothermal vents, to study the habitat specificity of genes. The compendium of ANME could be used to investigate the distributional pattern of ANME archaeal communities in global cold seeps and vertical niche partitioning. Together with their syntrophic SRB, the physiological basis of ANME-SRB interactions could also be explored.

### Code Availability

The present study did not use custom scripts to generate these datasets. The parameters and versions of all the bioinformatics tools used for the analysis were described in the Methods section.

## Acknowledgments

We thank all authors from the originating and submitting laboratories who contributed to the generation of the metagenomic sequence data and all persons who developed the software used in this study. The work was supported by the Scientific Research Foundation of Third Institute of Oceanography, MNR (No. 2022025 and No 2023022), State Key Laboratory of Marine Geology, Tongji University (No. MGK202303), and China Postdoctoral Science Foundation (2022M723709).

## Author contributions

X.D. designed this study. Y.H. and X.D. performed the analyses. C.Z. contributed to assembly of part of samples. X.D., Y.H. and Z.S. interpreted the data. Z.Z., Y.P., J.L., Q.J. and Q.L. contributed to the data collection. Y.H. and X.D. wrote the paper, with input from all other authors.

